# Mapping molecular determinants of Ca_V_2.2 inhibition by RGK proteins and homologs in *Xenopus* oocytes

**DOI:** 10.1101/2022.06.21.496996

**Authors:** Yehezkel Sasson, Suraj Subramaniam, Tal Buki, Lior Almagor, Orna Chomsky-Hecht, Moshe Katz, Henry Puhl, Stephen R. Ikeda, Nathan Dascal, Joel A. Hirsch

## Abstract

The Ca_V_1 and Ca_V_2 families of voltage-dependent calcium channels play a crucial role in neurotransmitter release, excitation-contraction and many other cellular processes. Comprised of the membrane pore-forming α_1_, intracellular β and extracellular α_2_δ subunits, these channels have been targets for pharmacological intervention for decades. Physiological functions of Ca_V_ channels are attenuated by either constitutively or transiently bounds proteins in the cellular environment. The RGK (Rad, Gem, Rem, and Rem2) G-protein family potently inhibits Ca_V_1 and Ca_V_2 function in heterologous expression systems. RGK proteins bind to Ca_V_β and inhibit channel localization and activity by forming a ternary complex with Ca_V_α_1_. Here, we evaluated the influence of RGK proteins on Ca_V_2.2 channels heterologously expressed in *Xenopus* oocytes. Both Gem and Rad showed no nucleotide dependency on its inhibitory function on Ca_V_2.2. The G-domain and C-terminus could inhibit the Ca_V_2.2 channel independently when co-expressed with channel subunits. Our results demonstrated that structural determinants in Gem, crucial for channel inhibition, lie within the 222-296 amino acid region containing both the partial G-domain and C-terminus as determined from chimeric Ca_V_β-Gem constructs. We expanded our mapping efforts and prepared various chimeras of *Drosophila melanogaster* (*Dm*) RGK sequences fused to Ca_V_β and showed that 22 residues in RGK2t and RGK3L C-terminal imparted complete Ca_V_2.2 inhibition. Point mutations in the *Dm*RGK C-terminus, conserved in mammalian RGK proteins, abrogated the Ca_V_2.2 inhibition to a significant extent, pointing to a hot region in the extreme C-terminus for inhibition of Ca_V_ channels. Since RGK homologs are now recognized as physiological modulators in β-adrenergic regulation of Ca_V_ channels, the relevance of this curious G-protein family deserves close examination.

## Introduction

The RGK mammalian gene family was discovered and cloned in the 1990s as transcriptionally regulated genes encoding small G-proteins. The founding four members (Rad, Gem, Rem, Rem2) all have a conserved architecture, encoding a central G-domain that binds guanosine nucleotides, flanked by a seventy residue N-terminus with little predicted structure and a fifty residue C-terminus (Kelly, 2005; Opatowsky et al., 2006; Yang and Colecraft, 2013). The last twenty residues in the C-terminus direct membrane localization but there is no evidence of any post-translational modification (Bilan et al., 1998; Heo et al., 2006; Splingard et al., 2007). In 2014, we reported that the RGK family was not limited to vertebrate animals but had homologs in the invertebrate world, hence members are also extant for the protostome (Puhl et al., 2014). The sequence motif which uniquely distinguishes RGK family genes and homologs is a conserved region of approximately eleven amino acids at the extreme C-terminus of the open reading frame.

RGK proteins are potent inhibitors of high voltage-activated calcium channels in heterologous systems. They are known to inhibit all Ca_V_1 and Ca_V_2 channels (Beguin et al., 2001; Finlin et al., 2003; Beguin et al., 2005a; Beguin et al., 2005b; Finlin et al., 2005; Beguin et al., 2006; Bannister et al., 2008; Fan et al., 2010; Xu et al., 2010) but have no effect on the low voltage-activated (Ca_V_3) channels (Finlin et al., 2003; Chen et al., 2005; Fan et al., 2010). Ca_V_1 and Ca_V_2 channels are multi-subunit complexes comprising the membrane pore-forming α1 subunit, an intracellular β subunit and an extracellular subunit, α2δ, that is glycolipid anchored to the plasma membrane (Dolphin, 2013, 2016). RGK proteins mediate their inhibitory effect via direct interaction with the Ca_V_β channel subunit (Beguin et al., 2001; Finlin et al., 2003; Ward et al., 2004; Beguin et al., 2005a; Chen et al., 2005; Andres et al., 2006; Correll et al., 2008b). Inhibition is not due to competitive association between Ca_V_α1 and RGK to Ca_V_β. Rather, ternary complex formation of Ca_V_α1, Ca_V_β and the RGK protein is essential for proper RGK function (Finlin et al., 2005; Yang et al., 2007; Correll et al., 2008b; Yang et al., 2010). Inhibition can be caused by both reduction of Ca_V_ channel cell surface density (Beguin et al., 2001; Beguin et al., 2005a; Beguin et al., 2006) and the inhibition of the I_Ca_ on the cell surface (Chen et al., 2005; Leyris et al., 2009; Fan et al., 2010; Yang et al., 2010; Fan et al., 2012; Yang et al., 2012). Recent studies demonstrate that calcium channel inhibition by RGK proteins is physiologically relevant as Rad plays a central role in adrenergic regulation of Ca_V_1 channels in cardiac myocytes (Ahern et al., 2019; Liu et al., 2020; Katz et al., 2021).

The structural determinants of RGK proteins important for Ca_V_ channel inhibition are thought to be located both in the G-domain and the C-terminus (Finlin et al., 2003; Chen et al., 2005; Beguin et al., 2006; Yang et al., 2007; Fan et al., 2012). While some studies show full inhibition can be achieved by the C-terminus alone (Leyris et al., 2009; Fan et al., 2012), others have shown that the G-domain, when sent to the membrane by a heterologous membrane localization signal, is able to impart the full inhibitory effect (Beguin et al., 2006). Together with these contradictory findings and inconsistencies, the ability of RGK proteins to work as molecular switches depending on their nucleotide binding state is also debated. While some studies show GTP-dependent inhibition (Ward et al., 2004; Beguin et al., 2005a; Beguin et al., 2005b; Beguin et al., 2006; Yang et al., 2010) others show no importance of nucleotide identity for RGK function (Bannister et al., 2008; Xu et al., 2010).

Here, we addressed two key questions. (*i*) Whether the nucleotide (GDP or GTP) bound to RGKs influences its function on Ca_V_ channel current inhibition? (*ii*) What are the precise RGK structural determinants involved in channel inhibition? Therefore, we engineered several constructs with RGKs and expressed them in a heterologous system, *Xenopus* oocytes, and employed the two-electrode voltage clamp (TEVC) approach. Our results show that both Gem and Rad show no nucleotide dependency for their inhibition of an effector, the Ca_V_2.2 channel. Employing a novel chimeric platform, we mapped determinants to residues 222-243 in the G-domain and residues 244-296 in the C-terminus of Gem. We also found that the conserved C-terminus of *Drosophila* RGK-like homologs, exhibited Ca_V_2.2 inhibition to a significant extent and point mutations in this region abrogated the inhibition.

## Materials and Methods

### DNA constructs

Rabbit genes encoding Ca_V_2.2 α1 (GenBank: D14157), Ca_V_β2b (ENA: L06110) and Ca_V_α2δ-1 (UniProt: P13806) were employed. All constructs included a T7 promoter sequence for *in vitro* RNA transcription. RNA transcripts included 5’ and 3’ untranslated sequences of *Xenopus* β-globin as mentioned previously (Dascal N., 1992). Human Gem (U10550), Rad (L24564) and H-Ras (P01112) cDNAs were amplified by PCR and inserted into pGEM-HJ between BamHI and XbaI restriction sites. Point mutations were created using overlapping PCR. The construction of Ras-Gem chimeras was performed by overlapping PCR. Likewise, Ca_V_β_core_-GS linker chimeras were prepared by overlapping PCR of Ca_V_β2b functional core (residues 1-424) with a linker, encoding 15 amino acids of Gly-Ser repetitions (GCGSGSGSGSGSGSGPR).Triple mutant Ca_V_β2b (1-424) (Ca_V_β_core_TM) was prepared by introduction of mutations (D244A, D320A, and D322A)(Opatowsky et al., 2003; Katz et al., 2021) into this functional core GS-rich linker cloned into the pGEM-HJ vector. Various lengths of Gem were cloned in tandem into the Ca_V_β_core_TM-GS linker C-terminus. *Drosophila melanogaster* RGK-like homologs RGK1 (AAF57577), 2t (AAF57577, amino acids 165-740), 3 (ABV53867, denoted as RGK3S in this study) and 3L (alternative splice variant of ABV53867), described previously (Puhl et al., 2014), were subcloned into pGEM-HJ vector. The various Ca_V_β_core_-RGK constructs were amplified by PCR and cloned into pGEM-HJ vector using Gibson assembly. Single- and double-point mutations in RGK2t and 3L were performed by standard overlapping PCR. All constructs were confirmed by DNA sequencing.

### In vitro RNA transcription

RNAs were prepared as previously described (Dascal N., 1992; Oz et al., 2017). Oocytes were injected with RNA two to four days before electrophysiological recording. In all experiments unless specified, Ca_V_β2b and α2δ-1 auxiliary subunits RNAs were co-injected with Ca_V_2.2 α1 in equal amounts by mass, ranging between 30-150 pg of each subunit. For Ca_V_β_core_-Gem and Ca_V_β_core_-RGK chimeras, the amount injected was also equal to Ca_V_2.2 α1 by mass. For Gem, Rad, Ras and Ras-Gem chimeras, the injected RNA amount was 2 ng per oocyte. Co-expression of RGK1, 2t, 3S and 3L in Fig. S4 was in the ratio of 2:1 (RGK: channel subunits) by mass. In Fig. S5A and B, concentrations of RNAs for Ca_V_β_core_-RGKs2t single point mutants were 2 ng each compared to 1 ng of Ca_V_2.2. Similarly, concentrations of RNAs for Ca_V_β_core_-RGKs3L single point mutants were 8 ng each.

### Electrophysiology

*Xenopus laevis* maintenance and oocyte preparation were as described (Shistik et al., 1998). Whole cell currents were recorded using the Gene Clamp 500B amplifier (Axon Instruments, Foster City, CA) using a two-electrode voltage clamp. Bath perfusion solution contained 50 mM NaOH, 2 mM KOH, 5 mM HEPES and 40 mM of Ba(OH)_2_ (titrated to pH 7.5 with methanesulfonic acid). Current-voltage (I-V) relations were measured with 15 ms pulses from a holding potential of -80 mV to test potentials of -50 mV to +50 mV in 10 mV steps. For each cell, the net currents were obtained by subtraction of the residual currents recorded with the same protocols after blocking the channels with 300 μM Cd^2+^. Data acquisition and analysis were performed using pCLAMP 9.0 software (Axon Instruments).

### *In-vitro* translation pulldown assay

Synthesis of S^35^-labled Ca_V_β chimeras was done by coupled in vitro transcription-translation (Promega (L1170)). Five nmole of purified 6xHisTag-MBP-AID was incubated with Ni-NTA beads in 30 mM HEPES pH 7.4, 200 mM NaCl, 5 mM MgCl_2_, 10% glycerol, 5 mM 2-mercaptoethanol, 1% triton and 20 mM imidazole for 1 hour at 4ºC. After incubation, beads were washed with the same solution for four times. Binding was initiated by adding 10 μl of the S^35^-labled Ca_V_β chimera to 250 μl beads solution for one hour on ice with constant mixing of the tube every 5 minutes. The Ni-NTA beads were than centrifuged at 1000 x*g* for one min and washed six times with one ml of the same solution containing 40 mM imidazole. Last wash was with 40 μl of the same solution and elution was carried out with 40 μl of the same solution containing 300 mM imidazole. Binding of the Ca_V_β -Gem chimeras was analyzed after SDS-PAGE by autoradiography.

### Statistical analysis

In all experiments, currents were normalized to the respective control (Ca_V_2.2 group), unless stated otherwise. Error bars are represented as mean ± standard error of the mean. Comparison between groups was done using one-way ANOVA for normally distributed data or Kruskal-Wallis ANOVA on ranks on skewed data. Holm-Sidak post hoc analysis was performed for normally distributed data and Dunn’s post hoc test, otherwise. Statistical analysis was performed using Sigmaplot 13 (Systat Software Inc. San Jose, CA, USA).

## Results

### Gem and Rad inhibit Ca_V_2.2 in a nucleotide-independent manner

Co-expression of WT Gem and Rad with Ca_V_2.2 exhibits almost complete inhibition of the Ba^2+^ current via Ca_V_2.2, I_Ba_ (Fig. 1). This result was shown previously for all RGK family members heterologously expressed in *Xenopus* oocytes and a diverse selection of human cell lines (Finlin et al., 2003; Beguin et al., 2005a; Chen et al., 2005; Finlin et al., 2005; Pang et al., 2010; Yang et al., 2010; Fan et al., 2012; Yang et al., 2012). We probed the importance of the nucleotide binding pocket and the nucleotide binding state by introduction of point mutations to residues in this region. The nucleotide binding affinities of several of these mutations have been determined previously (Sasson et al., 2011). Specifically, the Gem double mutant E83A, Q84P exhibits impaired GTP binding compared to the WT protein. Gem W133 is highly conserved across all RGK proteins, located in switch II of the G-domain. Finally, the Gem S89N mutant abolishes GTP binding and impairs GDP binding by two orders of magnitude. Moreover, the equivalent mutant in Rad, S105N, generates a protein that binds GDP exclusively. As seen in Fig. 1, both Gem/Rad WT and mutants displayed reduced I_Ba_ compared to the control, Ca_V_2.2 alone (P<0.001), whereas no significant differences in the I_Ba_ were observed between groups expressing Gem/Rad or their mutants, leading us to conclude that severely compromised nucleotide binding has no effect on RGK channel inhibition and demonstrating that nucleotide switching has no bearing on RGK’s effect on Ca_V_2 channel action.

**Fig. 1.**
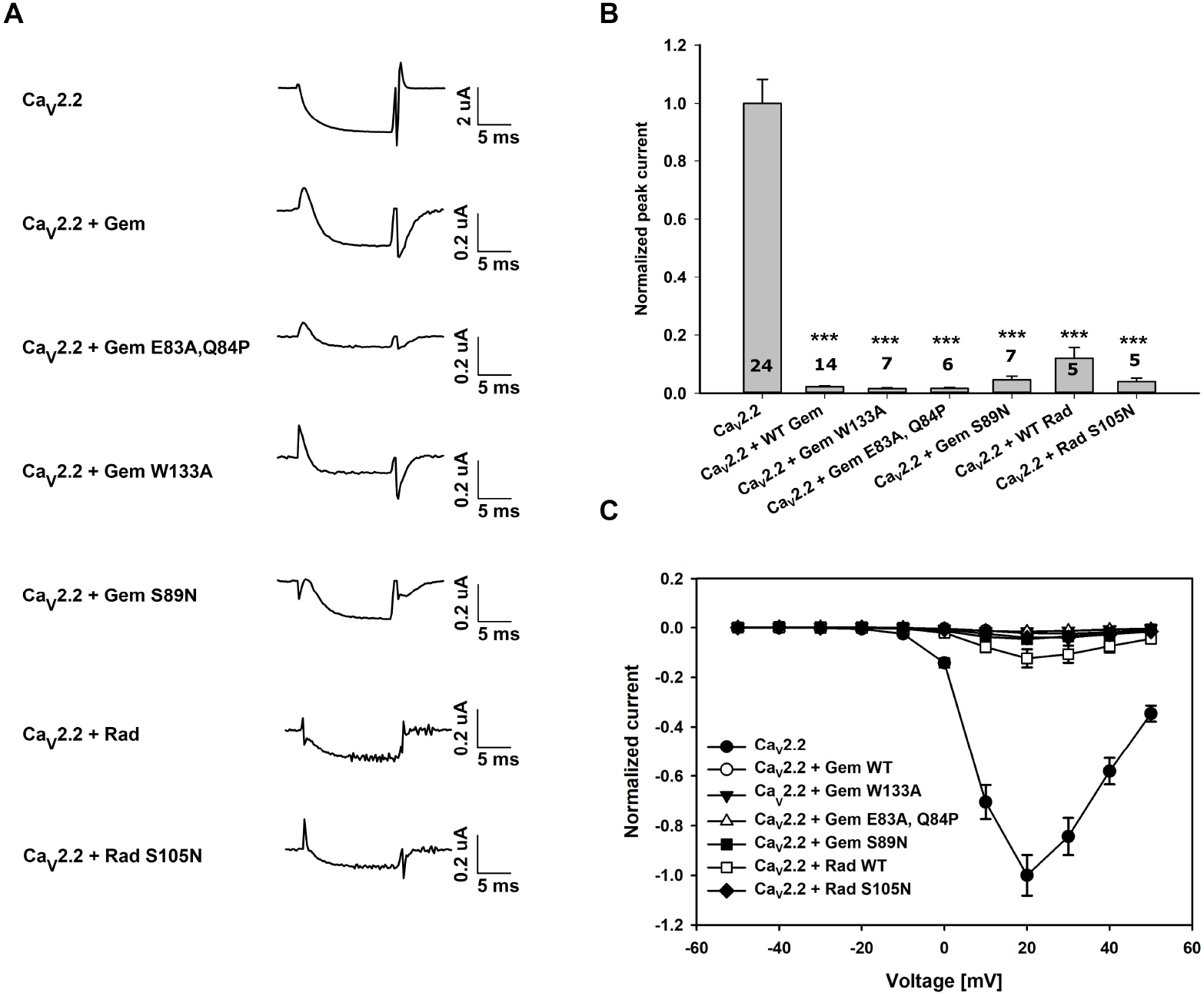
Ca_V_2.2 inhibition by WT Gem and Rad different point mutants. (A). Representative traces of I_Ba_^2+^ recorded during depolarization from -80 mV to + 20 mV applied to oocytes expressing Ca_V_2.2 + α_2_δ + β2b cRNAs and Gem or Rad cRNAs. (B) Representative traces of peak currents are shown in (A). All currents were normalized to the Ca_V_2.2 peak current which served as control. WT Gem and Rad and the mutants all showed almost complete inhibition of the measured I_Ba_ (C) I-V curves of the constructs mentioned in (A). (*) indicates statistical significance versus the control group. Statistics: One-Way ANOVA (P<0.001). The number of cells tested are indicated on the bar. *: P < 0.05; **: P < 0.01; ***: P < 0.001.

### The Gem C-terminus is a potent Ca_V_2.2 channel inhibitor

We next sought to clarify the importance of the Gem G-domain and C-terminus for Gem’s function. It remains an open question whether the G-domain contains intrinsic structural determinants important for inhibitory function or rather serves as a protein-protein interaction platform for a functional C-terminus. We examined this question in two independent ways. In the first approach, two chimeras of HRas and Gem were prepared: (*i*) a chimera composed of the HRas G-domain (residues 1-166) upstream of a Gem C-terminus (residues 244-296) and named RasG-Gem_CT_ and (*ii*) a chimera comprising the HRas G-domain (residues 1-143) upstream of Gem residues 222-296, named RasG-Gem_222-296_ (Fig. 2A). The latter chimera was created since this region of Gem (residues 222-296), expressed as a mini-protein, showed full I_Ba_ inhibition when tested in *Xenopus* oocytes (Leyris et al., 2009). HRas was selected as the chimeric G domain since the RGK proteins, as based on primary sequence analysis, belong to the Ras subfamily of the five subfamilies in the RAS superfamily (Colicelli, 2004), as we sought to best preserve the overall small G-protein architecture. Our working assumption designing these experiments was that HRas by itself does not associate with Ca_V_β, in contrast to the known association by RGK proteins. Significant inhibition of I_Ba_ by both chimeras was obtained in all groups compared to Ca_V_2.2 co-expressed with full length Ras (Ras_FL_) (Ca_V_2.2 + Ras_FL_; control) P<0.001 (Fig. 2C-D). Curiously, when Ca_V_2.2 and Ras_FL_ were co-expressed, a twenty percent larger I_Ba_ was observed compared to Ca_V_2.2 alone (P<0.001). Notably, experimental evidence of H-Ras-Ca_V_β2 interactions demonstrated recently (Servili et al., 2018) may play a role in these modestly elevated currents. Co-expression of RasG-Gem_CT_ and RasG-Gem_222-296_ with Ca_V_2.2 led to significantly smaller I_Ba_ as compared to control group (P<0.001) (Fig. 2C-D). However, a difference was also obtained between groups co-expressing RasG-Gem_CT_ versus RasG-Gem_222-296_ with Ca_V_2.2 (P=0.029) indicating that Gem_CT_ may possess the minimal necessary structural determinants required for Ca_V_2.2 channel inhibition. The fact that we observed partial inhibition leads us to conclude that membrane localization, due to the C-terminus (Heo et al., 2006), may provide some degree of affinity for the holo-channel without a direct Ca_V_β-Gem interaction. Alternatively, the Ras G-domain may enable direct interaction with Ca_V_β, albeit to a lesser degree than RGK proteins (Servili et al., 2018).

**Fig. 2.**
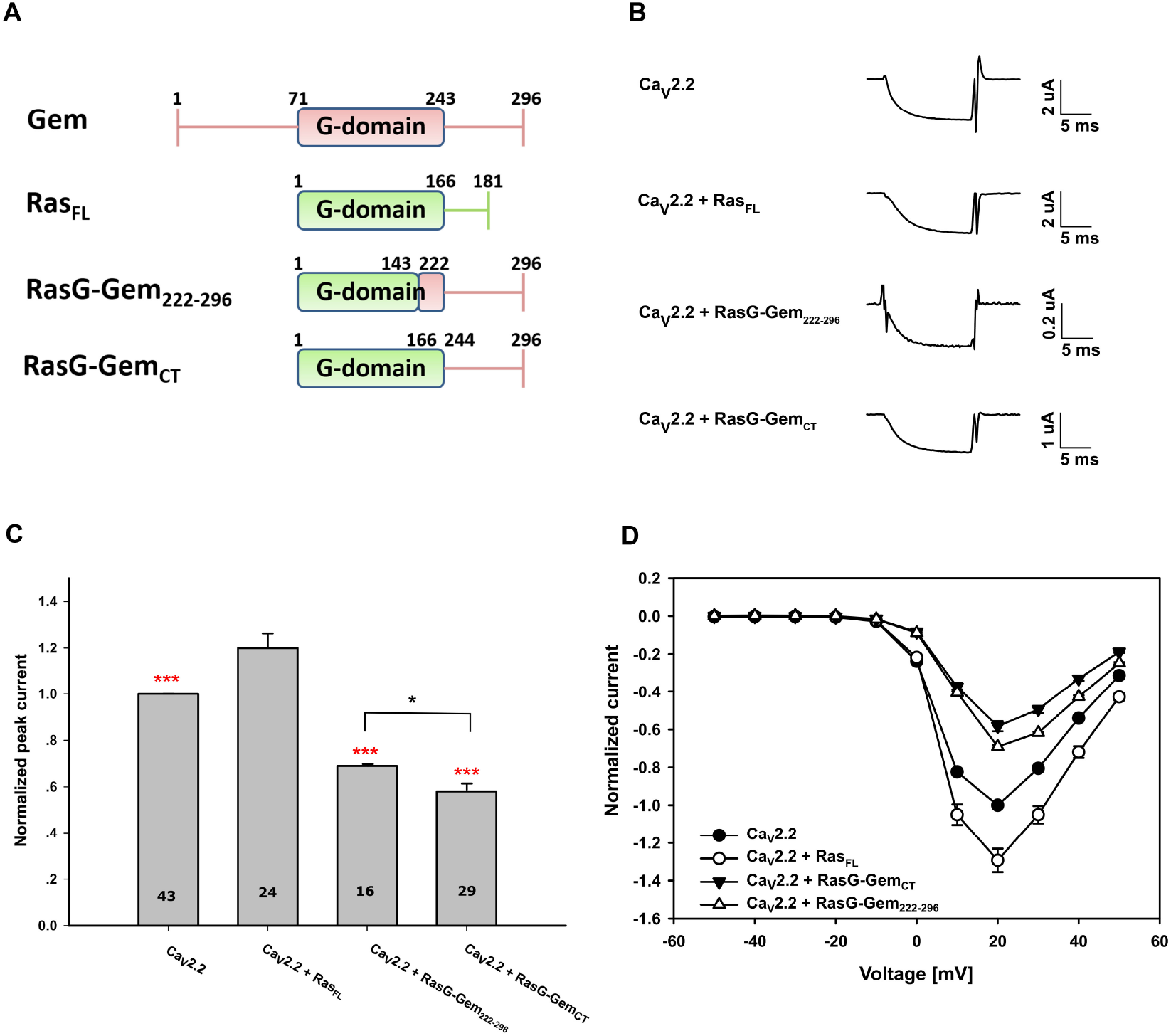
Ca_V_2.2 inhibition by Ras-Gem chimeras. (A) A schematic overview of the different constructs measured. (B) Representative traces of I_Ba_^2+^ recorded during depolarization from -80 mV to + 20 mV applied to oocytes expressing Ca_V_2.2 + α_2_δ + β2b cRNAs and cRNAs of different Ras-Gem chimeras. (C) Representative traces of peak currents of the constructs mentioned in (A). All currents were normalized to the Ca_V_2.2 peak current. Significant differences in I_Ba_ were obtained in groups. Red asterisk indicates significant differences between Ca_V_2.2+Ras_FL_ (control) and other groups. (D) I-V curves of the constructs mentioned in (A). Black asterisk indicates significant differences between the denoted groups. The number of cells tested are indicated on the bar. Statistics: One-Way ANOVA (P<0.001). *: P < 0.05; **: P < 0.01; ***: P < 0.001.

### Part of the Gem G-domain and C-terminus exhibit a combined inhibitory effect on Ca_V_2.2

The Ras-Gem chimeras demonstrate that the Gem C-terminus (residues 244-296) inhibits the Ca_V_ channel when co-expressed as an engineered G-protein. Next, we aimed to ascertain the crucial structural elements in the Gem G-domain and C-terminus crucial for Ca_V_2.2 channel inhibition with a different protein engineering strategy. To this end, we prepared fusion chimeras of the Ca_V_β functional core (residues 1-424) (Opatowsky et al., 2003) and diverse Gem constructs connected by a fifteen-residue Gly-Ser linker (Fig. 3A). These chimeras were designed to localize various Gem domains to the channel via fusion with Ca_V_β, testing their function without the requirement for a non-covalent Ca_V_β-Gem interaction. Variable lengths of Gem sequence were fused to the Ca_V_β functional core in the C-terminus rather than full length Ca_V_β to avoid having a ∼200 residue disordered region separating the two respective domains or proteins. Previously, it was shown that the Ca_V_β functional core is sufficient for proper function of this subunit (Opatowsky et al., 2003). No significant differences (P=0.138) in I_Ba_ were observed between full length Ca_V_β2b, Ca_V_β2b-core and Ca_V_β2b-core with the GS-linker on its C-terminus (Fig. S1 and (Katz et al., 2021)). Therefore, all Ca_V_β_core_- Gem chimeras were analyzed in comparison with the Ca_V_β functional core-GS linker construct. The Ca_V_β_core_-full length Gem (Ca_V_β_core_-Gem_FL_, Fig. 3) chimera showed complete reduction of the I_Ba_ (P<0.001) (Fig. 3C, extreme right histogram bar) indicating that the linker does not interfere with Gem’s inhibitory function. All the Ca_V_β_core_-Gem chimeras (Fig. 3C) exhibited a significant decrease in I_Ba_ (P<0.001) including Gem G-domain (residues 71-243) fused to Ca_V_β-core (Ca_V_β_core_-Gem_71-243_) (P=0.03) (Fig. 3B-D), when compared to the Ca_V_β_core_-GS linker control construct. Gem residues 222-243, which contains helix 5 of the G-domain, fused to Ca_V_β (Ca_V_β_core_-Gem_222-243_) and Gem residues 244-296 fused to Ca_V_β (Ca_V_β_core_-Gem_244-296_, Fig. 3A) showed significantly lower I_Ba_ compared to the Ca_V_β_core_-GS linker (Fig. 3B-D) indicating that both structural elements contribute to channel inhibition independently. However, a longer Gem sequence (residues 222-296) when fused to the Ca_V_β functional core (Ca_V_β_core_-Gem_222-296_, Fig. 3A), showed comparable inhibition of I_Ba_ to Ca_V_β_core_-Gem_FL_ (Fig. 3B-D), as a paired statistical test between these two constructs showed P=0.932. From these results we conclude that two primary structural elements impart Gem’s inhibitory function: *(i)* residues 222-243 and *(ii)* residues 243-296. Hence, when both these regions are included (residues 222-296), an additive inhibitory effect was observed with significantly smaller Ba^2+^ currents compared to chimeras bearing the individual elements, namely Ca_V_β_core_-Gem_222-243_ (P=0.037) and Ca_V_β_core_-Gem_244-296_ (P=0.031).

**Fig. 3.**
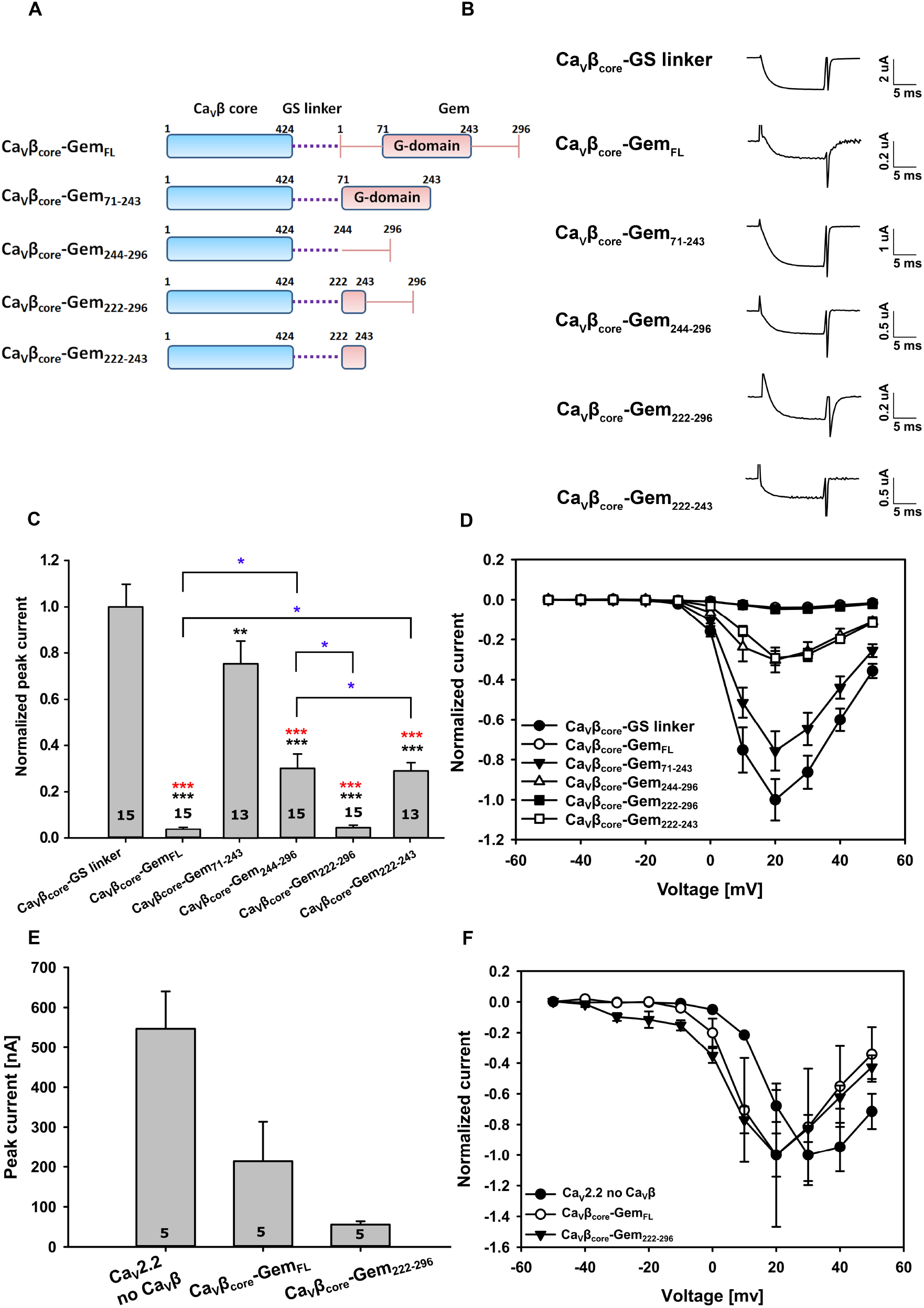
Ca_V_2.2 inhibition by Ca_V_β_core_-Gem chimeras. (A) A schematic overview of the different Ca_V_β_core_-Gem chimeras measured. (B) Representative traces of I_Ba_^2+^ recorded during depolarization from -80 mV to + 20 mV applied to oocytes expressing Ca_V_2.2 + α_2_δ cRNAs and cRNAs of different Ca_V_β-Gem chimeras. (C) Representative traces of peak currents of constructs mentioned in (A). Currents were normalized to the Ca_V_β_core_-GS linker peak current. Black asterisk indicates significant differences between the peak currents of all Ca_V_β_core_-Gem chimeras compared to Ca_V_β-GS linker group. Red asterisk indicates significant differences Ca_V_β_core_-Gem_71-243_ and other Ca_V_β-Gem chimeras. Blue asterisk indicates significant differences between the denoted groups. (D) I-V curves of the constructs mentioned in (A). (E) Averaged peak currents after 6 days. (F) I-V curves of the β-Gem chimeras compared to Ca_V_2.2 without Ca_V_β group after 6 days. Each I-V curve was normalized to its own peak current. Statistics: One-Way ANOVA (P<0.001). The number of cells tested are indicated in the histogram bar. *: P < 0.05; **: P < 0.01; ***: P < 0.001.

To ascertain whether the inhibition of I_Ba_ is caused by functional Gem in this novel chimeric framework or is simply a protein engineering artifact that gives rise to misfolded or non-functional Ca_V_β, we evaluated the shape of the I-V curve measured from these Ca_V_β_core_-Gem chimeras. As reported previously, presence of Ca_V_β causes a hyperpolarizing (leftward) shift in the I-V curve when co-expressed with Ca_V_2.1 (De Waard et al., 1994) and Ca_V_1.2 (Singer et al., 1991) as compared to oocytes lacking Ca_V_β subunit. I-V curves of Ca_V_β_core_-Gem_FL_ and Ca_V_β_core_-Gem_222-296_ were measured after six days in order to have large currents to assess I-V curves, as in the case of no Ca_V_β co-expression (Fig. 3F, filled circles). In addition, current amplitudes at +20 mV are presented in Fig. 3E. As seen in Fig. 3F, indeed co-expression of Ca_V_β_core_-Gem_FL_ and Ca_V_β_core_-Gem_222-296_ with Ca_V_2.2 causes a hyperpolarized shift as compared to Ca_V_2.2 without Ca_V_β subunit expression. These results demonstrate that in the Ca_V_β_core_-Gem chimeras, both Ca_V_β_core_ and Gem functioned.

### Localization of the Gem C-terminus to the holo-channel is sufficient for Gem inhibition

Previous reports demonstrate that the Ca_V_β-RGK interaction is crucial for RGK function on Ca_V_ channels, however, this interaction, characterized by charge-charge interactions, can be abolished by use of a mutant Ca_V_β bearing three point mutations, D244A, D320A, and D322A (hereafter, Ca_V_βTM) (Beguin et al., 2007; Yang et al., 2012). Our aim was to create a chimeric construct of Gem with this Ca_V_βTM backbone and assess the influence of the G-domain and C-terminal tail on Ca_V_2.2 inhibition when the Gem polypeptide is not bound to Ca_V_β in the usual three dimensional spatial configuration. Co-expressing Gem with Ca_V_β_core_TM that does not interact with Gem, resulted in the inability of Gem to inhibit the I_Ba_, as expected, while under similar conditions with Ca_V_β_core_, Gem almost completely abolished I_Ba_ (Fig. S2). When variable lengths of Gem were fused to this Ca_V_β_core_TM in the chimeric framework, the identical inhibitory properties of the different WT Ca_V_β_core_-Gem chimeras described above were retained (Fig. 4A-C, D). Significantly smaller I_Ba_ was observed in Ca_V_β_core_TM-Gem_FL_ and Ca_V_β_core_TM-Gem_222-296_, Ca_V_β_core_TM-Gem_244-296_ and Ca_V_β_core_TM-Gem_222-243_ (P<0.001) compared to Ca_V_β_core_TM-GS linker. However, no significant difference in I_Ba_^2+^ currents was detected between Ca_V_β_core_TM-Gem_71-243_ and Ca_V_β_core_TM-GS linker groups. Significant differences in the Ba^2+^ currents were observed between Ca_V_β_core_TM-Gem_71-243_ and other Ca_V_β_core_TM-Gem chimeras used (P<0.001). In conclusion, chimeric constructs with intact Gem C-termini (fused toCa_V_β_core_TM), remained unaffected by the triple mutation in the Ca_V_β_core_ and exhibited a large decrease in I_Ba_ thereby producing effective inhibition on the effector, the Ca_V_2.2 channel. In addition, the Gem G-domain (residues 71-243) which contains the residues required for the Ca_V_β_core_-Gem interaction, loses its inhibitory properties, observed in figure 3, when expressed as a chimera with the Ca_V_β_core_TM, despite its localization to the holo-channel via the GS linker.

**Fig. 4.**
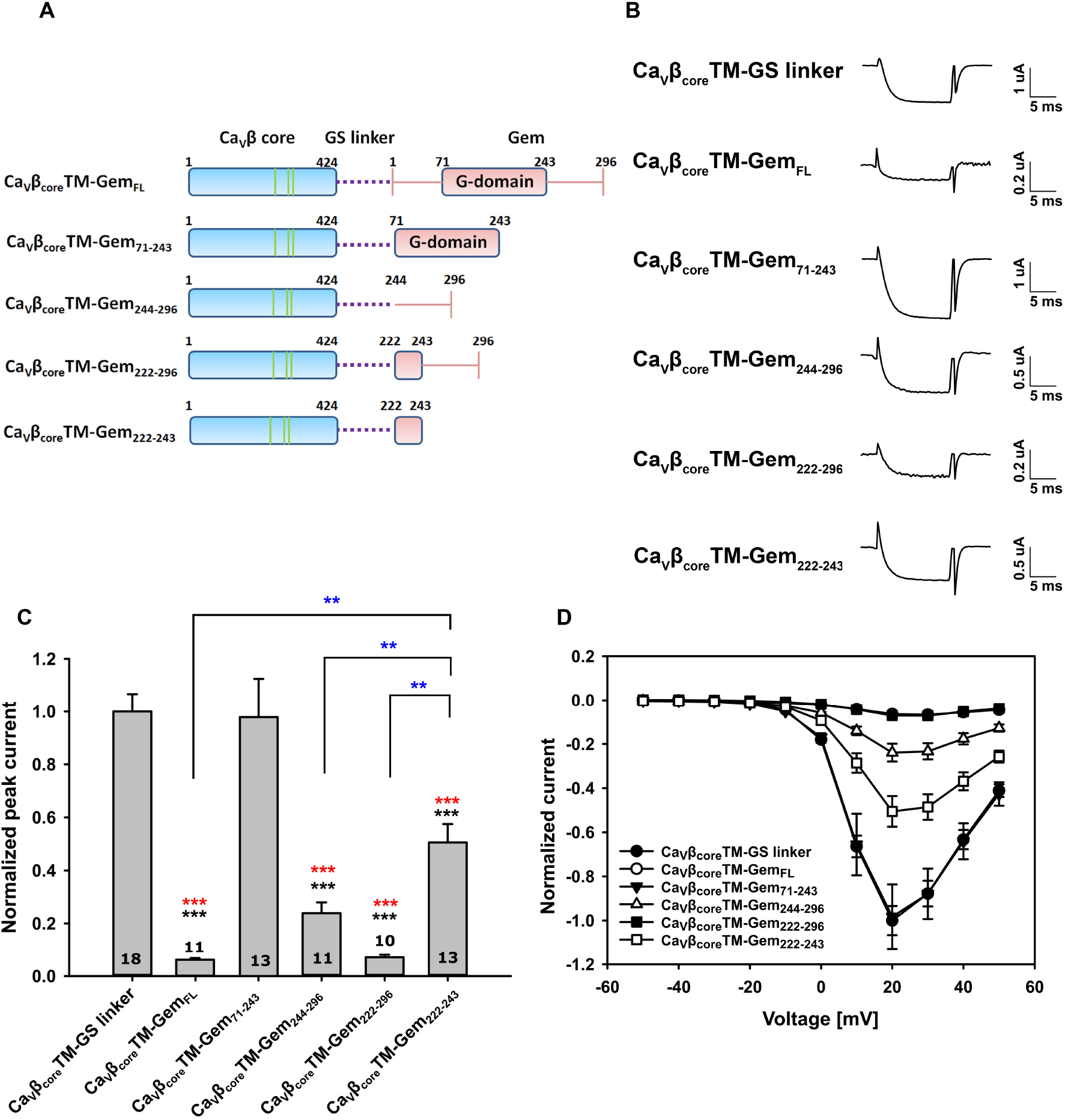
Ca_V_2.2 inhibition by Ca_V_β_core_TM-Gem chimeras. (A) A schematic overview of the different Ca_V_β_core_TM-Gem chimeras measured. Green lines represent three-point mutations in the Ca_V_β_core_TM (D244A, D320A & D322A) (B) Representative traces of I_Ba_^2+^ recorded during depolarization from - 80 mV to + 20 mV applied to oocytes expressing Ca_V_2.2 + α_2_δ cRNAs and cRNAs of different Ca_V_β_core_TM-Gem chimeras. (C) Averaged peak currents of the samples displayed in (A). Currents were normalized to the Ca_V_β_core_TM-GS linker peak current. Significant differences were observed between the peak currents of Ca_V_β_core_TM-GS linker shown by black asterisk and between Ca_V_β_core_TM-Gem_71-243_ shown by red asterisk compared to the other chimeras. The blue asterisk indicates significant difference between the denoted groups. No significant differences in the peak currents were observed between Ca_V_β_core_TM-Gem_71-243_ to Ca_V_β_core_TM-GS linker. (D) I-V curves of the constructs mentioned in (A). The peak current of Ca_V_2.2 + Ca_V_β_core_TM-GS linker was normalized to 1 and other currents were calculated relative to it. Statistics: One-Way ANOVA (P<0.001). The number of cells tested are indicated in the histogram bar. *: P < 0.05; **: P < 0.01; ***: P < 0.001.

The Ca_V_β-Gem chimeras were further validated for their ability to bind the pore-forming α1 subunit. To this end, pulldown experiments were conducted using purified HisTag-MBP-AID fusion proteins immobilized on Ni-NTA beads. *In vitro* translated S^35^ Ca_V_β_core_-Gem_FL_ and Ca_V_β_core_-Gem_222-296_, and their respective Ca_V_β_core_TM-GS linker versions, showed associations with AID as seen in Fig. S3. These associations were absent when HisTag-MBP alone was used as bait. This result demonstrates that the full Gem inhibitory effect was achieved without perturbing the Ca_V_β-AID interaction.

### RGK-like homologs from Drosophila melanogaster are potent inhibitors of Ca_V_2.2

Our previous report with the Ikeda group demonstrated that RGK-like proteins from (*Drosophila melanogaster, Dm*) significantly inhibit Ca_V_ channels in rat SCG neurons. A multiple sequence alignment presented in that study convincingly showed that the C-terminal ∼20 residues, especially the C-7 motif, remain highly conserved across arthropods to vertebrates (Puhl et al., 2014). Our aim was to first characterize the effects on Ca_V_2.2 by these RGK-like homologs from *Drosophila melanogaster* in *Xenopus* oocytes. Having validated their RGK-like function, namely Ca_V_ channel inhibition, we then sought to leverage the evolutionary divergence of the Protostome-Deuterostome split to reveal the minimal conserved structural elements encoding Ca_V_ channel inhibition. As reported previously, the predicted sequences of *Dm* RGK-like proteins are comprised of: *(i)* RGK1,*(ii)* RGK2t amino acids 165-740, *(iii)* RGK3 (denoted as RGK3S in this study) and *(iv)* RGK3L (splice variant of RGK3 with a ∼30 amino acid insertion after Q401) (Puhl et al., 2014). We co-expressed full length *Dm* RGKs 1, 2t, 3S and 3L with Ca_V_2.2 channel subunits in *Xenopus* oocytes. Similar to mammalian Gem and Rad, the *Dm* RGK-like homologs, RGK1, 2t and 3L indeed reduced I_Ba_ significantly (P<0.001) compared to Ca_V_2.2 expression alone (Fig. S4). RGK3S also showed significantly reduced (P=0.042) Ba^2+^ currents compared to Ca_V_2.2. However, this variant showed significantly larger currents in comparison to the oocyte group expressing RGK1 (P=0.002). This difference may be attributed to a short ∼30 amino acid region missing in RGK3S compared to RGK3L, which could possess some structural determinants crucial for channel inhibition (Fig. S4).

### Ca_V_β-DmRGK chimeras enable mapping of RGK determinants of channel inhibition

Having validated the functionality of the *Dm* RGK-like proteins in our heterologous system, we then used the chimeric approach described above and created three chimeras each for RGK2t and RGK3L, by fusing variable lengths of RGK2t and 3L sequences downstream of Ca_V_β_core_, separated by the GS linker (Fig. 5A). All six chimeras inhibited Ca_V_2.2 channels to varying degrees. Statistically significant differences were observed between the groups listed in Fig. 5C (P=0.002) and in Fig. 5E (P=0.001), respectively. In Fig. 5 C-D, the two RGK2t chimeras (Ca_V_β_core_-RGK2t_508-576_ and Ca_V_β_core_-RGK2t_537-576_) showed significantly smaller I_Ba_ (P=0.026 and P=0.002, respectively), than Ca_V_2.2 alone (control). The Ca_V_β_core_-RGK2t_FL,_ chimera showed complete inhibition of the channel to the point where we could not record measurable currents, despite several attempts which included titrating different concentrations of RNA (Fig. 5 C and D; P<0.001). Similarly, in Fig 5E-F, two RGK3L chimeras (Ca_V_β_core_-RGK3L_373-486_ and Ca_V_β_core_-RGK3L_447-486_) showed significantly smaller I_Ba_ compared to the Ca_V_2.2 alone (control) with P<0.001 and P=0.015, respectively. Chimera Ca_V_β_core_-RGK3L_FL_ clearly exhibited lower Ba^2+^ currents compared to Ca_V_2.2 alone, but lacked statistical significance (Fig. 5E-F). Overall, these results suggest that the structural determinants crucial for Ca_V_2.2 channel inhibition lie within the *Dm* RGK2t and RGK3L CT-tails (residues 537-576 and 447-486, respectively).

**Fig. 5.**
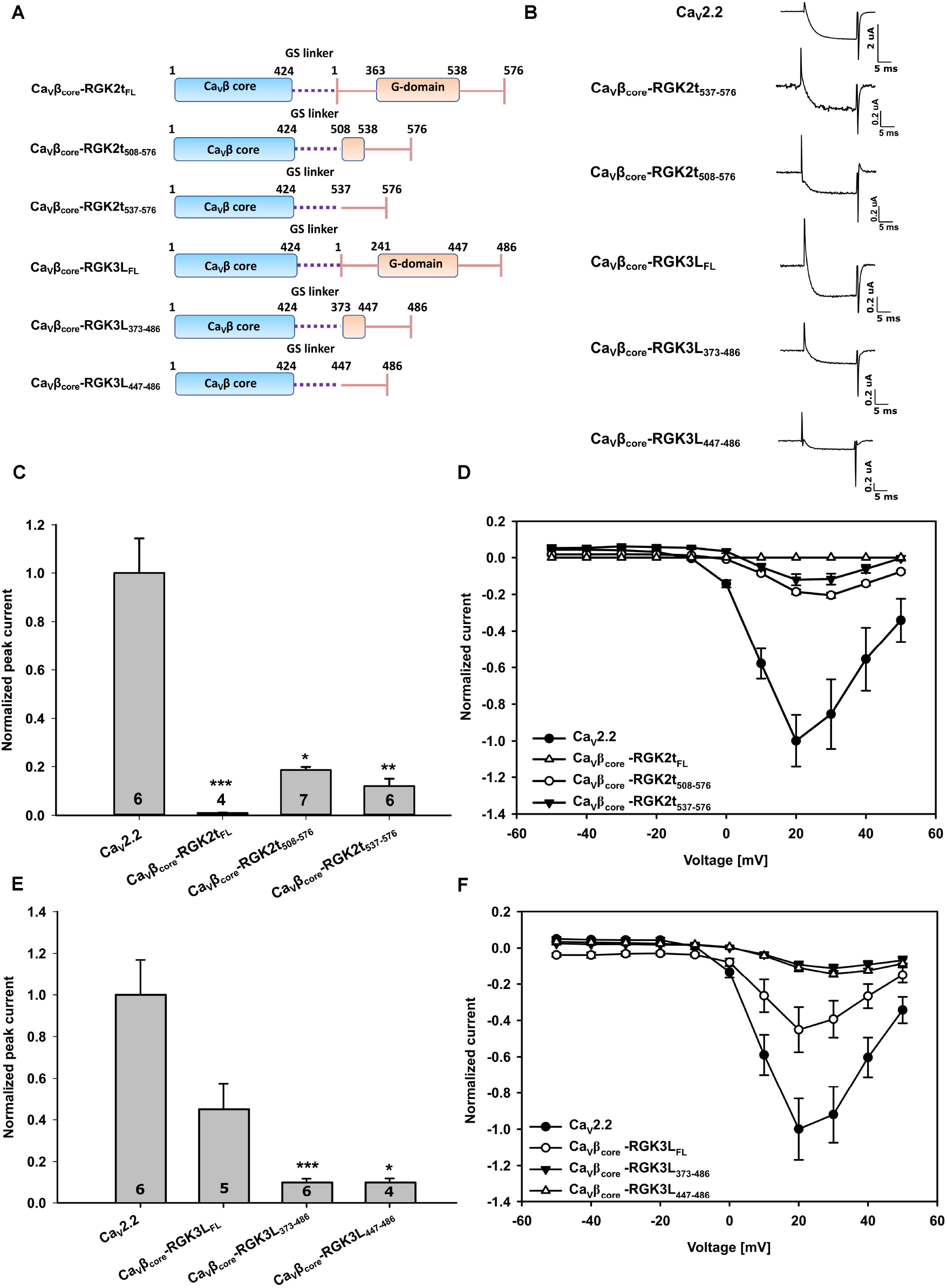
*Drosophila melanogaster* RGK chimeras are potent inhibitors of Ca_V_2.2. (A) A schematic overview of the different RGK2t and RGK3L chimeras fused to Ca_V_β_core_ are shown. (B) Representative traces of I_Ba_^2+^recorded during depolarization from -80 mV to + 20 mV applied to oocytes expressing Ca_V_2.2 + α_2_δ cRNAs and cRNAs of different Ca_V_β_core_-RGK2t and RGK3L chimeras. (C, E) Currents were normalized to the Ca_V_2.2 peak current (control) respectively. In (C), chimeras Ca_V_β_core_-RGK2t_508-576_ and Ca_V_β_core_-RGK2t_537-576_ showed significantly lower I_Ba_ compared to the control Ca_V_2.2 group (P=0.002). Similarly, in (E) chimeras Ca_V_β_core_-RGK3L_373-486_ and Ca_V_β_core_- RGK3L_447-486_ showed significantly lower I_Ba_ compared to the control Ca_V_2.2 group (P=0.001). (D and F) I-V curves of the constructs mentioned in (A). The peak current of Ca_V_2.2 was normalized to -1 and other currents were calculated relative to it. The number of cells tested are indicated in the bar. Statistics: Kruskal-Wallis One-Way ANOVA on ranks. (*) indicate significant differences versus the control group respectively. The number of cells tested are indicated in the histogram bar. *: P < 0.05; **: P < 0.01; ***: P < 0.001.

In an effort to resolve the RGK sequences responsible for inhibition, we used chimeras Ca_V_β_core_-RGK2t_537-576_ and Ca_V_β_core_-RGK3L_447-486_ as our structure-function probes, as they proved to be most effective in channel inhibition (Fig. 5). First, we incorporated single point mutations in the C-tail region (Fig. 6A and 5B). We chose to mutate cysteine 570 and leucine 573 in the RGK2t CT-tail and cysteine 480 and leucine 483 in the RGK3L CT-tail as these residues are either absolutely or very highly evolutionarily conserved (Puhl et al., 2014). As seen in Fig. 6C-D, single point substitutions with alanine in the RGK2t chimera continued to display inhibition of Ca_V_2.2. Substitutions of C570S or L573R gave the same result (Fig. S5A and B; not significant). Likewise, single point substitutions with alanine in the RGK3L chimera inhibited Ca_V_2.2 currents (Fig.6E-F), with parallel results for substitutions C480S or L483R (Fig. S5C and D; not significant). Next, we created double mutant versions of these chimeras, incorporating substitutions into both amino acids. Notably, while all chimeras showed significantly lower I_Ba_ compared to Ca_V_2.2 alone (P<0.001), the double point mutants significantly attenuated RGK inhibition of the Ca_V_2.2 channel (extreme right histogram bars in panels C and E, Fig. 6), by about fifty seventy percent, for RGK2t and 3L chimeras (P=0.006 and P=0.020, respectively). This result demonstrated that these conserved residues, cysteine and leucine are crucial for the RGK inhibitory effect on Ca_V_2.2 channels. Moreover, this loss of function phenotype required mutation of both residues.

**Fig. 6.**
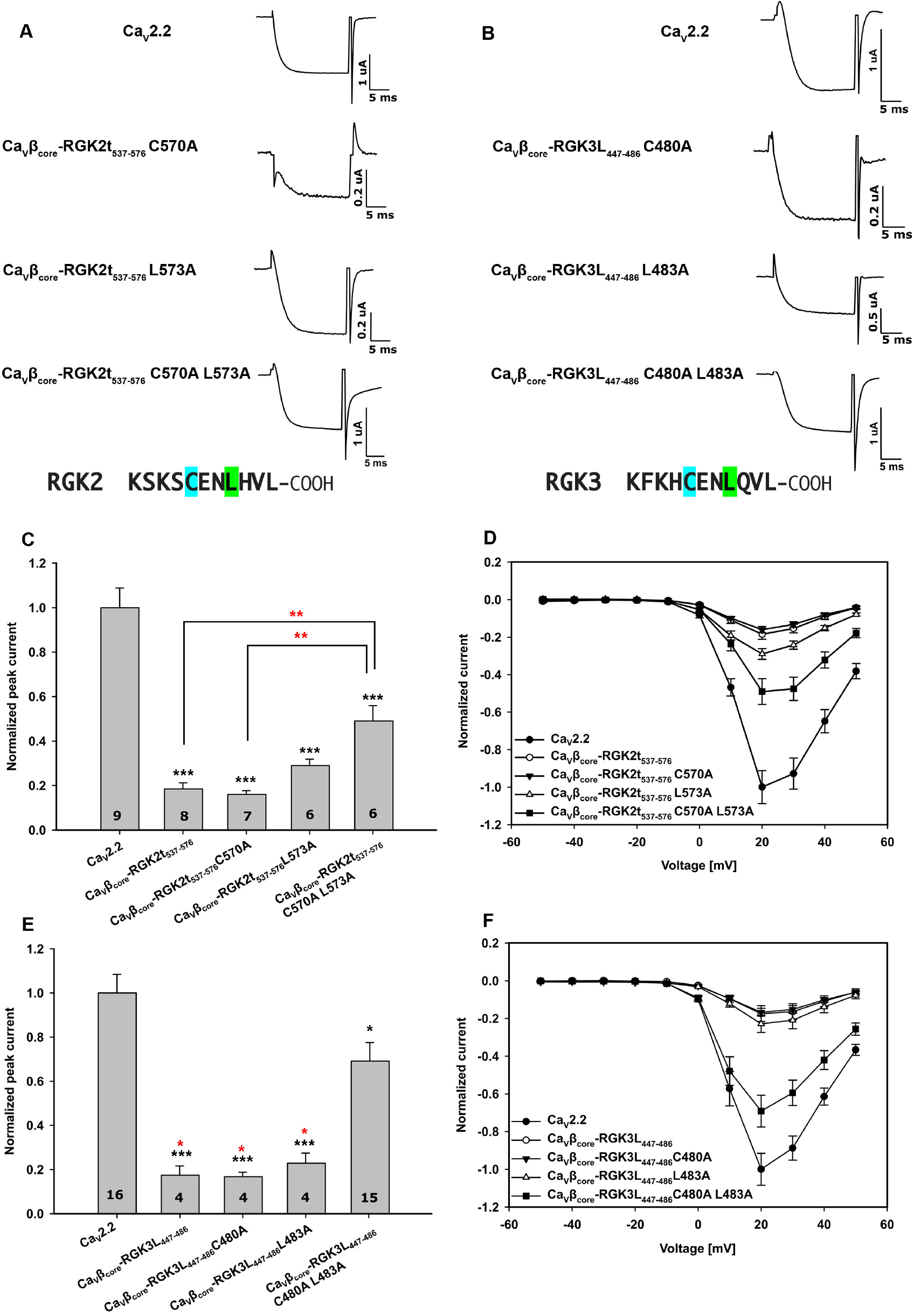
Double-point mutation in RGKs exhibited loss-of-function effect on Ca_V_2.2. (A, B) Representative traces of I_Ba_^2+^recorded during depolarization from -80 mV to + 20 mV applied to oocytes expressing Ca_V_2.2 + α_2_δ cRNAs and cRNAs of different Ca_V_β_core_-RGK2t and RGK3L chimeras with single- and double-point mutations. The last eleven C-terminal residues for RGK2 and RGK3 are shown. Highlighted in cyan and green are the cysteine and leucine residues that were mutated in the chimeras. (C, E) Currents were normalized to the Ca_V_2.2 peak current which served as the control group respectively. In (C), chimeras- Ca_V_β_core_-RGK2t_537-576_ along with single point mutants Ca_V_β_core_-RGK2t_537-576_ C570A and L573A and double point mutants Ca_V_β_core_-RGK2t_537-576_ C570A L573A showed significantly lower I_Ba_ compared to the control Ca_V_2.2 group. However, the double mutant showed significantly greater I_Ba_ than the Ca_V_β_core_-RGK2t_537-576_ and Ca_V_β_core_-RGK2t_537-576_ C570A chimeras. Black asterisk indicates significant differences versus the control group. Red asterisk indicates significant difference between the denoted groups. (D) I-V curves of the constructs mentioned in (A). Similarly, in (E) chimeras-Ca_V_β_core_-RGK3L_447-486_ along with single point mutants Ca_V_β_core_-RGK3L_447-486_ C480A and L483A and double point mutants Ca_V_β_core_-RGK3L_447-486_ C480A L483A showed significantly lower I_Ba_ compared to the control group. However, the double mutant showed significantly greater I_Ba_ than the Ca_V_β_core_-RGK3L_447-486_, Ca_V_β_core_-RGK3L_447-486_ C480A and Ca_V_β_core_-RGK3L_447-486_ L483A. Black asterisk indicates indicate significant differences versus the control group. Red asterisk indicates significant difference observed compared to Ca_V_β_core_-RGK3L_447-486_ C480A L483A and other groups. (F) I-V curves of the constructs of panel B. The peak current of Ca_V_2.2 was normalized to 1 and other currents were calculated relative to it. The number of cells tested are indicated in the bar. Statistics: One-Way ANOVA (P<0.001). The number of cells tested are indicated in the histogram bar. *: P < 0.05; **: P < 0.01; ***: P < 0.001.

## Discussion

The physiological role of RGK proteins in the cell has been enigmatic. Results from initial studies showed that RGK expression underwent transcriptional regulation, suggestive that they act as classical G-protein signaling molecules as based on sequence homology (Reynet and Kahn, 1993; Maguire et al., 1994; Finlin and Andres, 1997; Finlin et al., 2000). Strikingly, reports demonstrating genetic deletions of several RGK members were inconclusive in determining precise physiological function and relevance (Chang et al., 2007; Gunton et al., 2012; Magyar et al., 2012; Manning et al., 2013). Subsequent to their initial cloning, Beguin et al discovered the RGK:Ca_V_β interaction, revealing RGK Ca_V_ channel inhibition (Beguin et al., 2007); however, its physiological relevance remained undetermined. Recent studies have now shown *bona fide* physiological relevance for Rad as being a key player in the potentiation of Ca_V_1.2 channel currents during β-adrenergic regulation of cardiomyocytes and HEK cells, as mediated by cAMP and protein kinase A (Ahern et al., 2019; Liu et al., 2020). Our recent work extended this finding, where we showed involvement of Rad in both β1-or β2-mediated adrenergic regulation of Ca_V_1.2 in *Xenopus* oocytes (Katz et al., 2021). Like Rad, other RGK proteins may play similar physiological role(s) in β-adrenergic regulation of other tissues. Paradis et al., have shown Rem2-mediated inhibition of CaMKII concomitantly causes negative regulation of dendritic complexity (Ghiretti et al., 2013; Ghiretti et al., 2014). Surprisingly, direct regulation of calcium currents was not reported using RNAi-based methods on Rem2 in cultured neurons (Wang et al., 2011; Moore et al., 2013). Hence, according to the Paradis group, since Rem2 does not directly regulate inward Ca^2+^ currents, they propose that Rem 2 and other RGK proteins like Rem2 may instead be involved in orthogonal signaling pathways that remain poorly explored to date. Older studies of the cytoskeleton revealed that both Gem and Rad negatively regulate Rho kinase thereby playing a critical role in neurite remodeling (Ward et al., 2002). Notably, Rad has been reported to localize to the nucleus and affect transcription via nuclear factor κB inhibition (Hsiao et al., 2014).

Mounting evidence from this study and previous reports clearly suggests that the G-domain largely facilitates localization of the functional RGK C-terminus to the holo-channel, which is necessary for Ca_V_ inhibition. Our results align well with previous reports that demonstrated that deletion of the RGK C-terminus abrogates inhibitory function on Ca_V_ channels (Finlin et al., 2003; Chen et al., 2005; Yang et al., 2007; Leyris et al., 2009; Xu et al., 2010; Yang et al., 2010; Fan et al., 2012). Thus, the RGK C-terminus alone could serve as an effective Ca_V_ channel inhibitor when localized to the holo-channel either via G-domain or by fusion with Ca_V_β in our chimeric engineered constructs, imparting the inhibitory effect. As showed in previous reports, Gem’s inhibitory determinants map to its terminal 75 amino acids (Leyris et al., 2009; Fan et al., 2012). Here we report that the Gem C-terminal 75 amino acids possess two molecular determinants, each independently contributing to Ca_V_2.2 inhibition.

While some studies show GTP binding to be important for RGK function (Ward et al., 2004; Beguin et al., 2005a; Beguin et al., 2005b; Beguin et al., 2006; Xu et al., 2010), others show no contribution of GTP/GDP binding state to their function (Chen et al., 2005; Leyris et al., 2009). The electrophysiological work presented here, shows that guanine nucleotide binding to Gem or Rad has no influence on its inhibitory activity for its effector, the Ca_V_2.2 channel. Our results clearly indicate that while the G-domain could be critical in anchoring the RGK C-terminus to the holo-channel, the G-domain *per se* behaves as a pseudoGTPase since the protein-protein interaction is not dependent on conformation-induced nucleotide binding. Accordingly, a recent review (Stiegler and Boggon, 2020) classified RGKs as pseudoGTPases that play crucial role as signaling molecules within the cell. The classification of pseudoGTPases is an ongoing process as several less-well annotated proteins are now being classified under this group compared to other pseudoenzyme classes (Stiegler and Boggon, 2020). Careful biochemical studies by our group (Opatowsky et al., 2006; Sasson et al., 2011) failed to show any hydrolytic activity for either Gem or Rad, buttressing their identification as pseudoGTPases. Hence, we conclude that RGK proteins do not serve as paradigmatic G-protein molecular switches i.e., fluctuating between GTP-bound ‘on’ and GDP-bound ‘off’ state vis a vis their action on calcium channels (Correll et al., 2008a; Yang et al., 2010).

To elaborate a structure-function map of Gem sequences responsible for Ca_V_2.2 channel inhibition, we constructed Ras-Gem fusion chimeras. The rationale behind using HRas as the framework for chimeras was that HRas would behave in a structurally neutral manner when co-expressed with Ca_V_2.2 channel as RGKs are a subfamily of the Ras family (Colicelli, 2004). To our surprise, the results (Fig. 2C) showed that HRas may even enhance I_Ba_. This finding is consistent with results reported by other groups previously in several neuronal cell types (Hescheler et al., 1991; Hahnel et al., 1992; Pollock and Rane, 1996; Fitzgerald and Dolphin, 1997; Lei et al., 1998). The enhanced I_Ba_ may be attributed to the observed Ca_V_β:H-Ras interaction (Servili et al., 2018) responsible for depolarization-induced gene activation described recently. Despite the potentiation of I_Ba_ by HRas, our results clearly demonstrate that in the context of Ras-Gem chimeras, the Gem C-terminus inhibited I_Ba_ Ca_V_2.2 currents, pointing to the Gem C-tail as possessing a primary molecular determinant(s) responsible for channel inhibition. We conclude that the Ca_V_β:G-protein interaction has some level of conservation in the Ras family clade of small G-proteins.

We then engineered Ca_V_β_core_-Gem fusion chimeras in order to map the structural determinants that are necessary and sufficient for Gem-mediated Ca_V_2.2 channel inhibition. This novel chimeric fusion replaced the Ca_V_β-Gem non-covalent interaction, obviating the need for a specific motif that is responsible for Ca_V_β-RGK interaction (Beguin et al., 2007). Such a fusion also facilitates appropriate localization of the functional RGK unit (Gem) to the holo-channel. We assessed the functionality of two such chimeric Ca_V_βs by assessing the shape of the I-V curve using a control group lacking Ca_V_β co-expression. The chimeras (Fig. 3F) showed a hyperpolarized shift versus the control group owing to the presence of a functional Ca_V_β_core_ characterized previously (Singer et al., 1991; De Waard et al., 1994). Therefore, both Ca_V_β_core_ and Gem remained functional in such a chimeric arrangement, with the holo-channel localized to the plasma membrane. In an orthogonal approach, using pull downs with *in-vitro* translated Ca_V_β_core_-Gem_FL_ and Ca_V_β_core_-Gem_222-296_ and purified MBP-AID as a bait, we demonstrated that complexation of Ca_V_β_core_ with the I-II linker and the AID specifically remained uncompromised in these chimeras. The chimeric Ca_V_β-Gem constructs show that Gem_222-296_ possesses two determinants, each independently contributing to Ca_V_2.2 inhibition. Our results align well with a previous report where structural determinants in Gem responsible for inhibiting the Ca_V_2.1 channel were mapped to the twelve amino acids in the C-terminus and may involve L241/R242/R243 amino acids in the Ca_V_β core (Fan et al., 2012).

Subsequently, we characterized chimeric constructs bearing a triple mutation (TM) in Ca_V_β_core_ and fused variable lengths of Gem to it, as done with wild-type Ca_V_β_core_. The rationale for this experimental series was to ascertain whether the native quaternary structure comprised of α_1B_, Gem and Ca_V_β is required for the positioning of Gem on the holo-channel in Gem-mediated inhibition. Our results show that the TM mutant chimeras remained functional, inhibiting channel currents. Chimeras with intact Gem C-termini showed maximal decrease in Ba^2+^ currents. Hence, chimeras-Ca_V_β_core_- Gem_FL_, Ca_V_β_core_-Gem_222-296_ and Ca_V_β_core_-Gem_244-296_ possess the principal molecular determinant(s) involved in reducing I_Ba_ Ca_V_2.2 currents but, importantly, do not need to be oriented by Ca_V_β in a specific spatial arrangement to affect channel activity with the following caveat. The partial inhibitory effect observed with the G-domain (residues 71-243) vanishes when in a chimeric arrangement with Ca_V_β_core_-TM as opposed to when fused with Ca_V_β_core_ (Fig. 4C vs Fig. 3C, respectively). We believe that this specific loss of partial inhibition may be due in part to improper positioning within the holo-channel. On the other hand, the three mutations of Ca_V_β_core_-TM disrupt the Ca_V_β_core_ GuK domain-Gem G-domain interface (Beguin et al., 2007), and presumably are also disrupted in the Ca_V_β_core_TM-Gem_71-243_ chimera. Taken together, results from the chimeric approach (Ca_V_β_core_TM-Gem) emphasize the importance of Gem C-termini in Gem-mediated inhibition. We posit that Gem’s membrane localization, in part, is mediated by interaction with Ca_V_β, and in turn Ca_V_β’s position within the holo-channel.

The development of Ca_V_β chimeras described in this study represents a novel platform for investigating Ca_V_ structure-function. Here, we have employed it for studying RGK function. In a complementary manner, Colecraft and coworkers [reviewed in (Colecraft, 2020)] fused PKCγ-C1 to various cytosolic proteins, namely Ca_V_β, CaM, and Rem. These engineered molecules behave has calcium channel blockers when activated with phorbol ester in HEK 293 cells. Contrary to their approach, our chimeric constructs have fused RGK elements fused to Ca_V_β_core_, which uses the poly-basic motif in the RGK C-terminus for membrane localization rather than the C1 domain. While it cannot be excluded that our Ca_V_β-RGK chimera may behave akin to the C1 fusion protein in the context of membrane anchoring and calcium channel inhibition, the RGK effect may be reversible via phosphorylation (Katz et al., 2021; Liu et al 2020), unlike C1-fused RGK proteins whose phorbol ester-dependent membrane association is practically speaking irreversible.

We previously reported that RGK-like proteins from *Drosophila melanogaster* inhibited I_Ca_ in rat superior cervical ganglion neurons (Puhl et al., 2014). In the current study, we corroborated this finding, showing that *Dm*RGKs could inhibit mammalian Ca_V_2.2 channels in the heterologous oocyte system. Having confirmed their functionality in oocytes, we then used a parallel molecular chimeric approach wherein variable lengths of RGK2t and 3L were fused to functional Ca_V_β_core_ to characterize the channel inhibition and to further map the determinants in these *Dm*RGKs. Our goal was to decipher molecular determinants in these evolutionarily distant *Dm*RGK homologs that nonetheless possess the conserved C-termini and exhibit similar functional activities as mammalian RGKs. In this manner, we leverage evolution to identify the minimal necessary but sufficient structure-function determinants required for channel inhibition. Analogous to highly conserved enzyme active-sites (Jack et al., 2016), the functional RGK C-terminus has likely remained conserved during evolution (Puhl et al., 2014) due to strong selection pressure. Our findings reveal that both *Dm*RGK C-tails potently inhibit the channel independently without the involvement of a G-domain, unlike Gem (residues 222-243) that possesses determinants in both the G-domain and C-tail. In the case of *Dm*RGK single point mutants, the identity of the substitution i.e., cysteine to serine/alanine or leucine to arginine/alanine in both RGK2t and 3L chimeras (Fig. S5) did not markedly attenuate the inhibitory function. This finding implies that substitution of cysteine with a polar or non-polar residue or substitution of leucine with a positively charged or non-polar residue do not by themselves disrupt the C-tail’s interaction with its target. C-tail double-point mutants of these highly conserved amino acids (Puhl et al., 2014), indeed markedly weakened the inhibitory effect but did not abrogate it, implying that other RGK residues in both Ca_V_β_core_ RGK2t_537-576_ C570A L573A and Ca_V_β_core_ RGK3L_447-486_ C480A L483A constructs impart inhibition to some degree. We posit that the C-terminal region constitutes a “hot region,” constituting a dominant peptidic segment [reviewed in (London et al., 2013; Ozdemir et al., 2018)] mediating plasma membrane or protein interaction. Notably, the invariant ‘cysteine’ within the C-terminal seven amino acid sequence is absolutely conserved across all RGK proteins (Puhl et al., 2014). To date, no lipid modification of this cysteine has been reported. Further, in a comprehensive structural bioinformatic study of cysteine positions based upon structural databases, cysteines have “the highest tendency to be found in crucially important regions of proteins” and do not serve as generic hydrophobic or hydrophilic residues, unlike other amino acids (Marino and Gladyshev, 2010).

In framing these findings, we propose that the generic RGK protein behaves in a manner similar to channel toxins. Like toxins, RGK proteins inhibit channel activity and are soluble factors. In other words, RGKs may serve as endogenous, cell autonomous intracellular factors that inhibit calcium channels, as opposed to toxins which are exogenous factors that evolved as part of physiological defense systems. Another molecular precedent might be phospholamban, a cellular endogenous membrane protein that acts to inhibit the calcium pump, SERCA, in a reversible manner. Toxins primarily function via two known mechanisms: *(i)* pore blockers, that bind to the outer vestibule and block the ion flow; *(ii)* gating modifiers, interact with a channel region and alter channel conformation whilst opening or inactivating and influence the gating mechanism (Kalia et al., 2015). ω-conotoxin GVIA inhibits Ca_V_2 channels, occluding the Ca_Vα1_2.2 pore thereby preventing Ca^2+^ influx (McCleskey et al., 1987). But, unlike ω-conotoxin GVIA, that selectively binds to Ca_V_2.2 in a virtually irreversible manner, the effect of RGKs can be reversed (Liu et al., 2020; Katz et al., 2021) by their phosphorylation, releasing them from the holo-channel. Therefore, in this framework, the RGK in the Ca_V_β:RGK chimeric molecule, might occlude the cytoplasmic opening of Ca_Vα1_ pore by inserting itself, analogous to the ‘ball and chain’ model for Shaker channels (Armstrong, 1981) concomitantly decreasing the open probability of the channel. It remains quite possible that RGK proteins exhibit multiple modalities to regulate Ca_V_ channels: *(a)* directly, by physical occlusion of the Ca_V_α1 pore; *(b)* indirectly, by interacting with Ca_V_β subunit, restricting its movement in the holo-channel (Yang et al., 2010; Yang and Colecraft, 2013).

## Acknowledgments

This work was supported by Israel Science Foundation grants 1519/12, 1500/16, 2780/20 to JAH. SS was supported in part by a scholarship from the Prajs–Drimmer Institute at Tel Aviv University. We would like to acknowledge Louise M Silverman for her assistance in recordings. German-Israeli Science Foundation (GIF grant I-1452-203.13/2018) to N.D. and the Gessner Fund grant to M.K.

## Author contributions

Yehezhel Sasson, Suraj Subramaniam, Lior Almagor, Tal Buki, Orna Chomsky-Hecht, Moshe Katz performed the experiments; Yehezhel Sasson, Suraj Subramaniam, Lior Almagor, Tal Buki, Joel Hirsch and Nathan Dascal analyzed the data. Henry Puhl III and Steve Ikeda contributed reagents. Yehezhel Sasson, Suraj Subramaniam and Joel Hirsch wrote the paper. All authors read and approve the manuscript.

