## Supplementary information for "Mapping molecular determinants of Ca_V_2.2 inhibition by RGK proteins and homologs in *Xenopus* oocytes"

[1]Dept of Biochemistry and Molecular Biology
School of Neurobiology, Biochemistry, and Biophysics
George S. Wise Faculty of Life Sciences

[2]Dept of Physiology and Pharmacology
Sackler Faculty of Medicine

[3]Sagol School of Neuroscience
Tel Aviv University
Ramat Aviv 6997810
Israel

[4]Laboratory of Molecular Physiology, Section on Transmitter Signaling
National Institute on Alcohol Abuse and Alcoholism
National Institutes of Health
Rockville, MD
USA

*Authors contributed equally

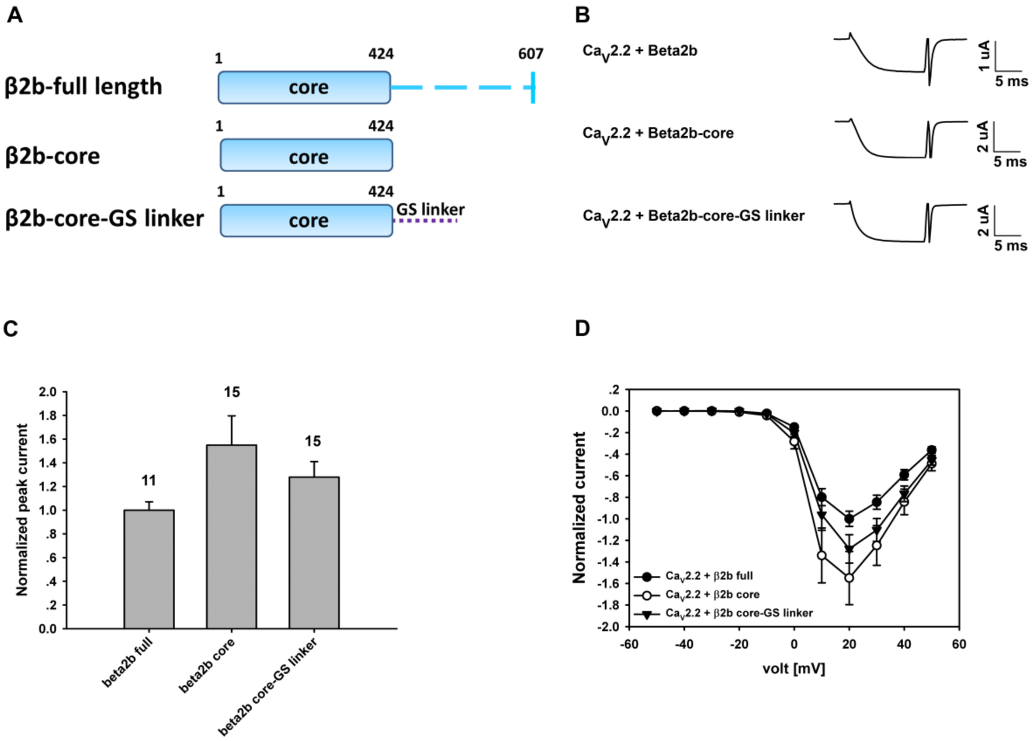


**Fig. S1. The GS linker has no influence on the $I_{Ba}^{2+}$ measured for $Ca_V2.2$** (A) A schematic overview of the different constructs measured. (B) Representative traces of $I_{Ba}^{2+}$ recorded during depolarization from -80 mV to + 20 mV applied to oocytes expressing $Ca_V2.2$ + $\alpha_2\delta$ cRNAs and cRNAs of $Ca_V\beta2b$, $Ca_V\beta2b$ core or $Ca_V\beta2b$ core-GS linker. (C) Averaged peak currents of the samples displayed in (A). Currents were normalized to the $Ca_V\beta2b$ full peak current. (D) I-V curves of the constructs mentioned in (A). No statistically significant differences were observed between the groups. Statistics: One-Way ANOVA (P=0.138). The number of cells tested is indicated above the bar.

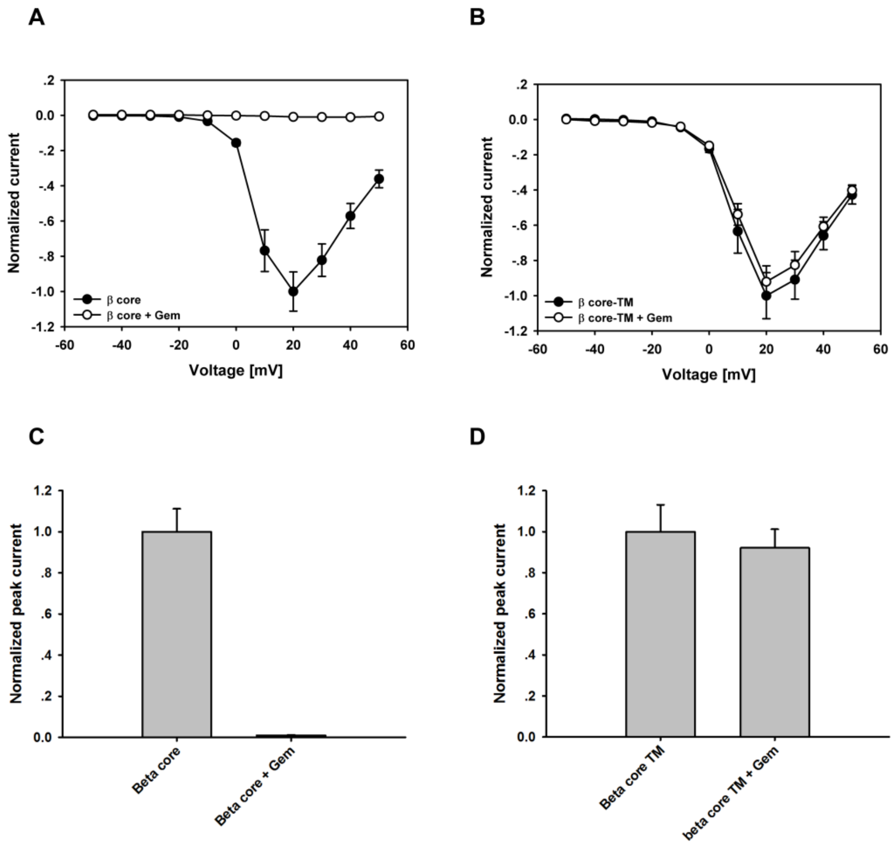


**Fig. S2. $Ca_V$2.2 inhibition by Gem is dependent on $Ca_V\beta$-Gem interaction.** (A) Averaged peak currents of $Ca_V$2.2 with or without Gem in the presence of $Ca_V\beta$2b (1-424) (B). Averaged peak currents of $Ca_V$2.2 with or without Gem in the presence of $Ca_V\beta$2bTM (1-424) ($Ca_V\beta$ with three-point mutations D244A, D320A & D322A, resulting in a Gem binding deficient $Ca_V\beta$). For both panels, currents were normalized to the current in the absence of Gem. (C) & (D) The I-V curves of the peak currents shown in (A) & (B), respectively. For panels A and C, n=8 oocytes while for panels B and D, n=12 oocytes.

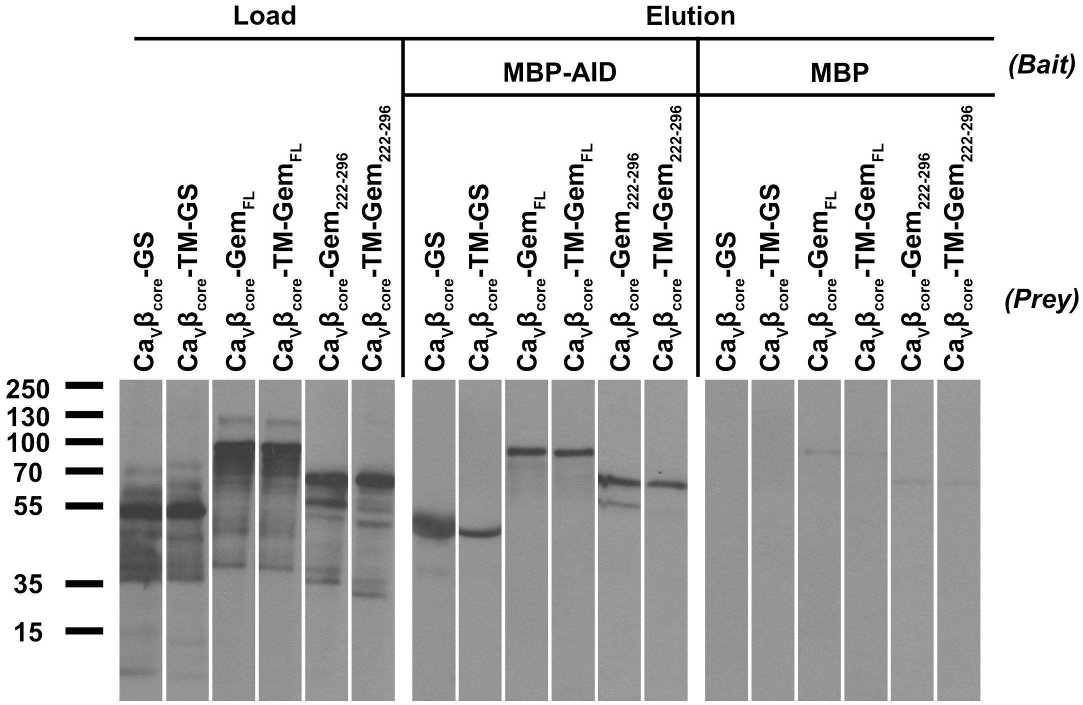


**Fig. S3. $Ca_V\beta$ chimeras do not disrupt the $Ca_V\beta$-$Ca_V\alpha 1$ interaction mediated by the AID.** Pull-down assays of in vitro translated $Ca_V\beta$2b-Gem chimeras. The left side (Load) of the autoradiogram shows the input $Ca_V\beta$2b-Gem chimeras constructs. The right side (Elution) shows the pulled down $Ca_V\beta$2b-Gem chimeras with recombinant VDCC MBP-AID eluted from amylose beads with maltose. MBP was used as negative control bait.

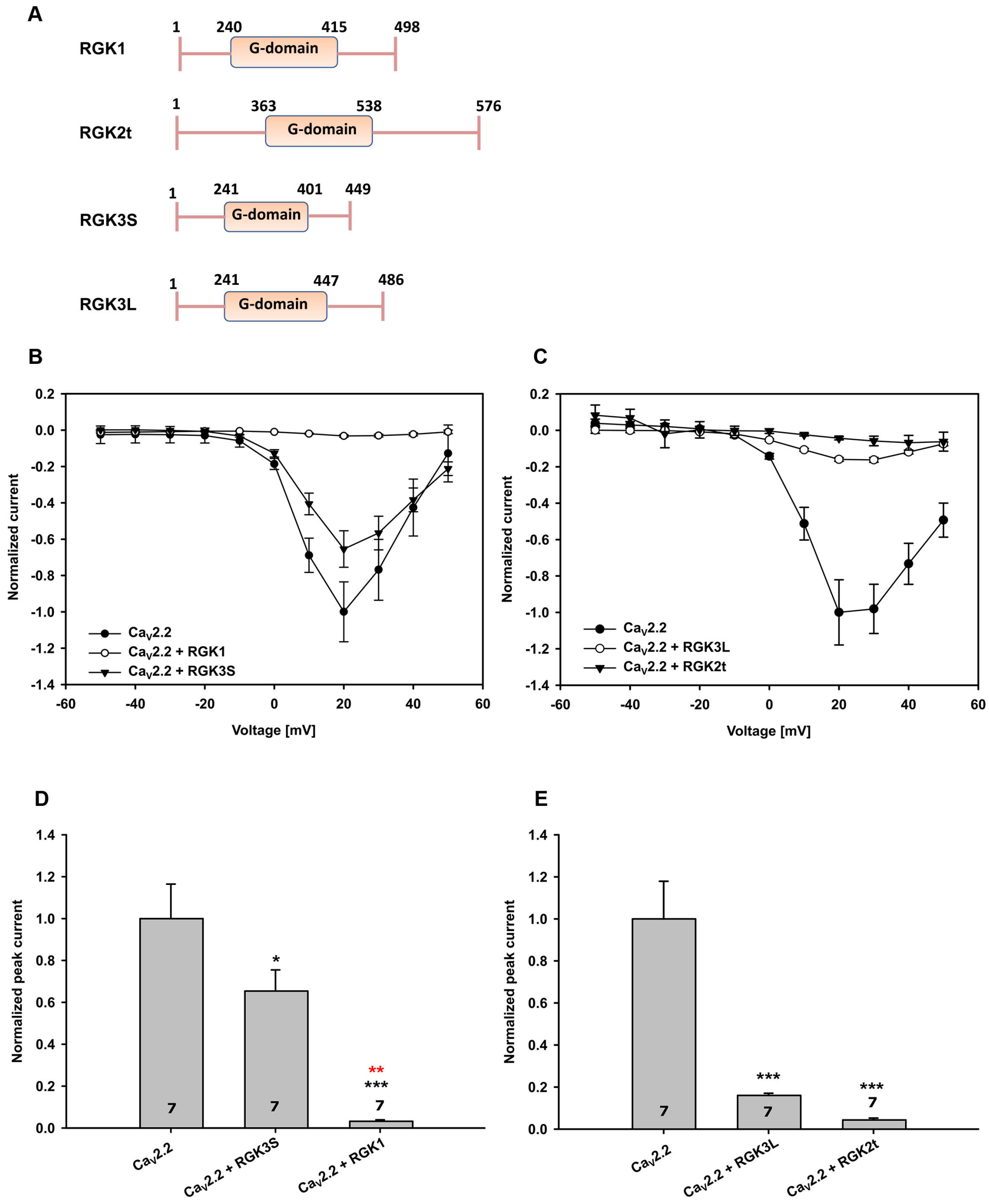


**Fig. S4. Full-length DmRGK1, 2t, 3L and 3S potently inhibit $Ca_V2.2$.**

(A) A schematic overview of the different constructs measured (B) I-V curves of $Ca_V2.2$ co-expressed with different full-length DmRGKs. RGKs1, 2t, 3L and 3S were co-expressed in the ratio of 1:2 (channel subunits: RGK). (C) All currents were normalized to the $Ca_V2.2$ peak current in (C) and (D). Co-expression of RGK3S and RGK1 with $Ca_V2.2$ showed significantly lower $I_{Ba}$. (*) indicates significant differences compared to the control $Ca_V2.2$ group. (*) indicates significant differences compared to $Ca_V2.2$+RGK3S. $Ca_V2.2$+RGK1 group showed significantly lower $I_{Ba}$ than $Ca_V2.2$ + RGK3S (D) Co-expression of RGK3L and RGK2t with $Ca_V2.2$ showed significantly lower $I_{Ba}$. (*) indicates significant differences compared to the control $Ca_V2.2$ group. Statistics: One-Way ANOVA (P<0.001). The number of cells tested are indicated on the bar. Notes: *P<0.05 **P<0.01 ***P<0.001

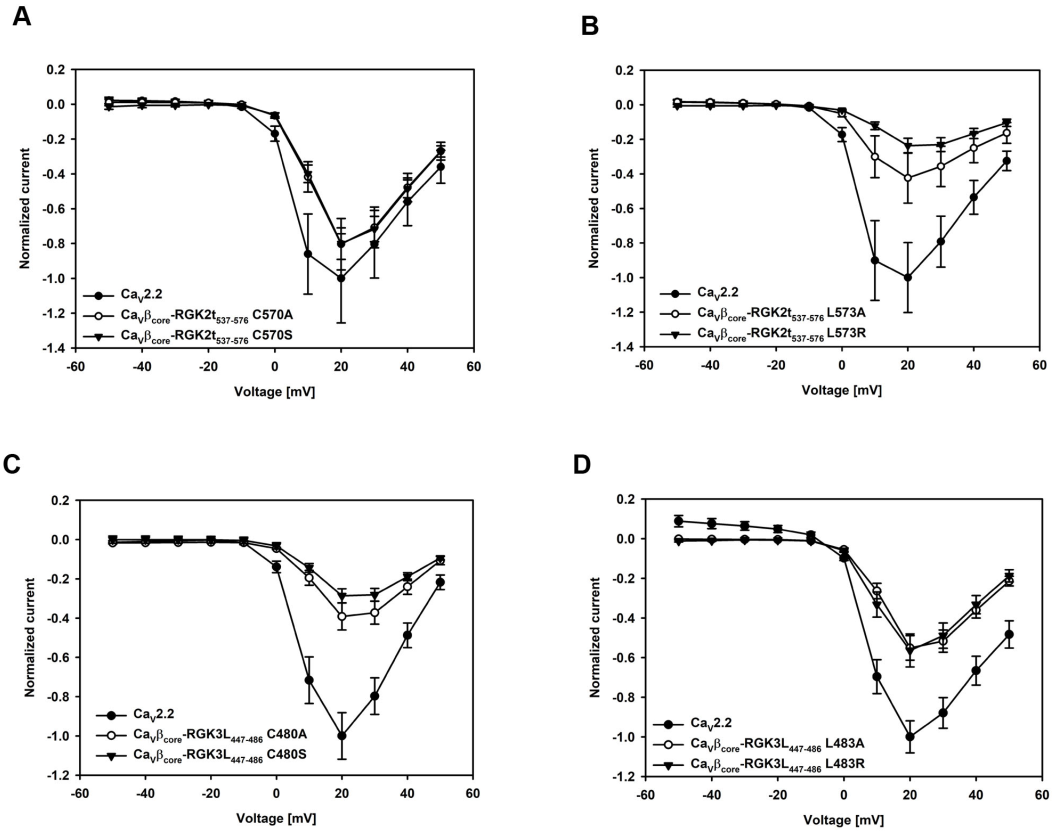


**Fig. S5. Single point mutations in the C-tail of RGK2t and 3L exhibited similar extent inhibition.** (A and B) Point mutations in the c-termini of constructs described in the panel of $Ca_V\beta_{core}$-RGK2t showed no-significant differences in $I_{Ba}$ compared to each other. (C and D) Point mutations in the C-termini of constructs described in the panel of $Ca_V\beta_{core}$-RGK3L showed no-significant differences in $I_{Ba}$ compared to each other.
